# Continuous chromatin state feature annotation of the human epigenome

**DOI:** 10.1101/473017

**Authors:** Bowen Chen, Neda Shokraneh Kenari, Maxwell W Libbrecht

## Abstract

Semi-automated genome annotation (SAGA) methods are widely used to understand genome activity and gene regulation. These methods take as input a set of sequencing-based assays of epigenomic activity (such as ChIP-seq measurements of histone modification and transcription factor binding), and output an annotation of the genome that assigns a chromatin state label to each genomic position. Existing SAGA methods have several limitations caused by the discrete annotation framework: such annotations cannot easily represent varying strengths of genomic elements, and they cannot easily represent combinatorial elements that simultaneously exhibit multiple types of activity. To remedy these limitations, we propose an annotation strategy that instead outputs a vector of chromatin state features at each position rather than a single discrete label. Continuous modeling is common in other fields, such as in topic modeling of text documents. We propose a method, epigenome-ssm, that uses a Kalman filter state space model to efficiently annotate the genome with chromatin state features. We show that chromatin state features from epigenome-ssm are more useful for several downstream applications than both continuous and discrete alternatives, including their ability to identify expressed genes and enhancers. Therefore, we expect that these continuous chromatin state features will be valuable reference annotations to be used in visualization and downstream analysis.

## 1 Introduction

Sequencing-based genomics assays can measure many types of genomic biochemical activity, including transcription factor binding, chromatin accessibility, transcription, and histone modifications. Data from sequencing-based genomics assays is now available from hundreds of human cellular conditions, including varying tissues, individuals, disease states, and drug perturbations.

Semi-automated genome annotation (SAGA) methods are widely used to understand genome activity and gene regulation. These algorithms take as input a collection of sequencing-based genomics data sets from a particular tissue. They output an annotation of the genome that assigns a label to each genomic position. They are semi-automated in the sense that they discover categories of activity (such as promoters, enhancers, genes, etc) without any prior knowledge of known genomic elements and a human interprets these categories, similar to a clustering algorithm. Many SAGA methods have been proposed, including HMMSeg [3], ChromHMM [6, 7], Segway [9] and others. See below for a comprehensive review of previous work.

All existing SAGA methods output a discrete annotation that assigns a single label to each position. This discrete annotation strategy has several limitations. First, discrete annotations cannot represent the strength of genomic elements. Variation among genomic elements in intensity or frequency of activity of cells in the sample is captured in variation in the intensity of the associated marks. Such variation is lost if all such elements are assigned the same label. In practice, SAGA methods often output several labels corresponding to the same type of activity with different strengths, such as “Promoter” and “WeakPromoter” [10, 6]. Second, a discrete annotation cannot represent combinatorial elements that simultaneously exhibit multiple types of activity. To model combinatorial activity, a discrete annotation must use a separate label to represent each pair (or triplet etc) of activity types. For example, intronic enhancers usually exhibit marks of both transcription and regulation. However, representing all possible combinations of activity types with discrete labels would require a number of labels that grows exponentially in the number of activity types.

In this work, we propose a continuous genome annotation strategy. That is, our method outputs a vector of real-valued *chromatin state features* for each genomic position, where each chromatin state feature putatively represents a different type of activity. Continuous chromatin state features have a number of benefits over discrete labels. First, chromatin state features preserve the underlying continuous nature of the input signal tracks, so they preserve more of the information present in the raw data. Second, in contrast to discrete labels, continuous features can easily capture the strength of a given element. Third, chromatin state features can easily handle positions with combinatorial activity by assigning a high weight to multiple features. Fourth, chromatin state features lend themselves to expressive visualizations because they project complex data sets onto a small number of dimensions that can be shown in a plot. For these reasons, in other fields, continuous modeling is often preferred over discrete. For example, the widely-used method of topic modeling for text documents assigns a continuous weight to each of a number of categories (such as “sports” or “politics”) for each document [12].

In this work we explore the utility of chromatin state feature annotation. We propose several measures of the quality of a chromatin state feature annotation and we compare the performance of several alternative methods according to these quality measures. We propose a Kalman filter state space model for this problem, epigenome-ssm, that produces the highest-quality continuous annotations of the methods we compared.

## 2 Related work

Many methods have been proposed for discrete chromatin state label annotation [3, 6, 9, 13, 22, 17, 26, 2]. The primary model used by these methods is the hidden Markov model (HMM). An HMM is a probabilistic model that assumes that there is a latent (unknown) chromatin state label at each position, and that the observed genomics data sets are generated as a function of this label. An HMM assumes that the label at position *i* depends only on the label at position *i* − 1. Later work extended this basic approach in a number of ways. First, there are three methods for modeling the input genomics data: one can binarize the data and model 0/1 values with a Bernoulli distribution [6], use a continuous measure of signal strength such as fold enrichment over control modelled with a Gaussian distribution [9, 22], or model raw read counts with a negative Binomial distribution [17]. Second, some methods [9] use statistical marginalization to handle unmappable regions. Third, several strategies exist for modelling segment lengths or for producing annotations on multiple length scales [9, 13]. Finally, several strategies have been proposed to guide the choice of the number of labels [22, 26, 24, 2].

A related class of joint annotation methods aim to improve epigenome annotations by simultaneously annotating many cell types and sharing position-specific information between the annotations [24, 25, 1, 15, 14]. Such joint annotations can be more accurate, but have the drawback that they may mask differences among cell types. We do not consider the joint annotation task in this work, but adapting the continuous annotation approach to this task is a promising direction for future work.

Another related task aims to take data from all available cell types as input to produce a single cell-type-agnostic (as opposed to cell-type-specific) annotation, a task known as “stacked” annotation. Several methods have been proposed to produce a discrete [10, 16] and continuous [5, 21] cell-type-agnostic annotations. However, these methods do not apply to the cell-type-specific case, so we do not compare to these methods below.

## 3 Methods

### 3.1 Reference genome

We performed all analysis using the human reference genome hg19.

To improve computational efficiency, following previous work [6, 16], we divided the genome into 200 bp bins and performed all analysis at the bin level. Note that we chose 200 bp to facilitate comparison to previous work; we recommend a finer resolution (e.g. 50 or 100 bp) in practice.

Following previous work [9], to train our model, we used just the ENCODE Pilot regions (http://hgdownload.soe.ucsc.edu/goldenPath/hg19/encodeDCC/referenceSequences/encodePilotRegions.hg19.bed), which cover about 1% of the human genome. We also removed the ENCODE blacklist regions (https://www.encodeproject.org/annotations/ENCSR636HFF/) from all analysis.

### 3.2 ChlP-seq histone modification data sets

We downloaded ChIP-seq histone modification data sets from the Roadmap Epigenomics data portal [11]. See [11] for a full description of the data processing pipeline. Briefly, reads are mapped to the reference genome, shifted and extended according to the fragment length and compared to an input control. We represented the signal at a given position as the negative log P-value of the observed read count compared to input, according to a Poisson model [11]. To reduce the influence of large outliers, following previous work [9], we transformed ChIP-seq values using the transformation arcinsh 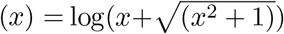. We used data from five histone modifications (H3K4me3, H3K4me1, H3K36me3, H3K27me3, H3K9me3) in the human lymphoblastoid cell line GM12878 (https://egg2.wustl.edu/roadmap/data/byFileType/signal/consolidated/macs2signal/pval/).

### 3.3 State space model

We developed a Kalman filter state space model [4] for annotating the genome with chromatin state features. This model takes as input a vector of *E* observed genomics data sets for each position, *y_g_* ∈ ℝ*^E^*, for *g* ∈ 1…*G*. This model assumes that at position *g* there is a latent vector *α_g_* ∈ ℝ^*M*^ that encodes the chromatin state features of that position. It assumes that the observed data vector at that position *y_g_* is generated as a linear function of *α_g_* plus Gaussian noise,

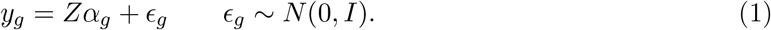

It further assumes that the latent vector *α_g+1_* is generated as a linear function of *α_g_* plus Gaussian noise,

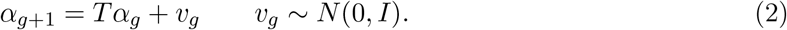

To learn the SSM model, we use the EM algorithm to maximize the log likelihood of the model as a function of its parameters, *Z* ∈ ℝ*^E×M^* and *T* ∈ ℝ*^M×M^* (Algorithm 1). Briefly, this algorithm alternates two steps, the E step and the M step. In the E step, we hold *Z* and *T* fixed and use a message-passing algorithm to efficiently estimate *α_1:g_* and compute sufficient statistics for updates to *Z* and *T*. In the M step, we use these sufficient statistics to update *Z* and *T*. In Section 4.1 below, we also consider a simpler algorithm which performs the E step independently by holding *Z,T,α_1:g−1_* and *α_g+1:_G* fixed and updating *α_g_*, then iterating for all *g*. We initialized *Z* ~ Uniform(0,1)^*E×M*^ and *T = I_M_*.

To limit the model’s capacity to overfit and its sensitivity to local optima, we additionally add several *L*_2_ regularization terms to the optimization’s objective function *J*(*Z, T*), which encourage *Z* and *T* to have small values:

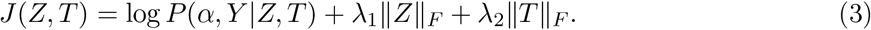

We parallelized the E step over a cluster by dividing the genome into windows, computing sufficient statistics from each window on worker nodes, and combining the sufficient statistics to compute updates to *Z* and *T* on a master node.

### 3.4 Hidden Markov model (HMM)

The most common model used in previously-proposed genome annotation is the hidden Markov model (HMM). The HMM assumes that at position *g* there is a latent state *x_g_* ∈ {1…*k*} that represents the chromatin state label of that position. It assumes that the observed data is generated as function of the latent state *x_g_* (see next paragraph). It further assumes that the state at position g depends just on the state at position *g* − 1, and that state *l* transitions to state *l*’ with probability *φ_ℓ,ℓ’_*

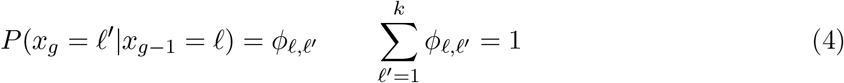

We considered two versions of the HMM model, which take as input continuous or discrete data respectively. As described in Section 2, some existing methods take continuous input [3, 9] while others take discrete input [8, 6]. In the continuous input case, as with the state space model, the continuous-input HMM takes as input a vector of *m* observed genomics data sets for each position, *y_g_* ∈ ℝ*^m^*, for *g* ∈ 1…*T*. It assumes that there is a mean vector associated with each state *μ_k_*, and that the observed data vector equals *μ_k_* plus Gaussian noise

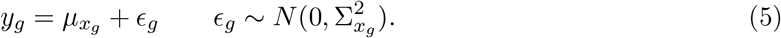

In the discrete input case, input data is thresholded into binary values such that the input data at position *g* is represented by a binary vector *ȳ_g_* ∈ {0, 1}^*m*^. The discrete-input HMM assumes that the observed data is generated as a multivariate Bernoulli distribution. That is, for track *i* at position *g*,

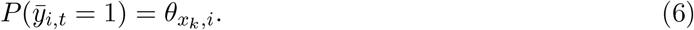

To learn the HMM model, we use the EM algorithm to maximize the log likelihood of the model as a function of the model’s parameters: *μ*_1:*k*_, Σ_1:*k*_ and *ϕ*_1:*k*,1:*k*_ for the continuous-input model, and *θ*_1:*m*,1:*k*_ and *ϕ*_1:*k*;1:*k*_ for the discrete-input model.

We consider two possible ways of representing the output of the HMM model. Existing methods generally output a discrete value the inferred state *x_g_* as the chromatin state label at position *g*. However, an HMM can be re-purposed to output continuous chromatin state features by defining the chromatin state feature *ℓ* at position *g* as *α_ℓ,t_* = *P*(*x_g_* = *ℓ*).

In summary, we have two choices for input (continuous input with a Gaussian distribution or discrete input with a Bernoulli distribution) and two choices for output (output continuous chromatin state features or discrete chromatin state labels), giving us four HMM variants: HMMgaus-con, HMMgaus-dis, HMMber-con and HMMber-dis. We used the Python package pomegranate [20] for all training and inference of HMM models.

### 3.5 Alternative models

We additionally tried three other alternative models. First, we tried simply using the raw input tracks as the chromatin state features. To produce a chromatin feature annotation with *k* features, we randomly selected a subset of *k* of the input tracks. Second, we tried principle component analysis (PCA) [18] and non-negative matrix factorization (NMF) [23]. Both of methods each take as input an input vector *y_g_* ∈ ℝ^*m*^ and output a vector of features *α_g_* ∈ ℝ^*k*^ to optimize a likelihood function. They differ from the SSM and HMM methods in that PCA and NMF treat each position independently, without considering the genome coordinate. NMF differs from PCA in that it outputs non-negative values *α_i,t_* ≥ 0. We used implementations of PCA and NMF from scikit-learn [19].

### 3.6 Existing reference annotation

For comparison, we downloaded the annotation results of ChromHMM from the Roadmap Epige-nomics data portal http://egg2.wustl.edu/roadmap/web_portal/chr_state_learning.html#core_15state, IDEAS http://bx.psu.edu/~yuzhang/Roadmap_ideas/test1.116.bb and Segway https://hoffmanlab.org/proj/segway/2012/segway_gm12878.bed.gz.

### 3.7 Gene expression evaluation

Following previous work [25], we evaluated an annotation according to the strength of correlation between the labels at a promoter of a given gene and that gene’s expression. we downloaded RNA-seq data from the Roadmap Epigenomics data portal. (http://egg2.wustl.edu/roadmap/data/byDataType/rna/expression/57epigenomes.RPKM.pc.gz)

We used a linear regression model to evaluate the degree to which annotations at a promoter region are predictive of gene expression. We trained two types of evaludation models: region-specific models and a whole-gene model. For the region-specific models, we divided each gene into 10 evenly-spaced bins. We also defined a bin for each 1 kb interval out to 5 kb upstream of the gene’s transcription start site (TSS) and 5 kb downstream of the gene’s transcription termination site (TTS), for a total of 20 bins.

For each bin, we trained a linear regression model. As the regression feature vector, we used the average feature vector in the respective bin. For discrete annotations, we used a one-hot encoding such that the feature vector has a 1 in the position corresponding to the label and 0’s elsewhere. As the regression response value, we used the RNA-seq RPKM (reads per kb per million mapped reads) values. As with ChIP-seq data, we transformed RPKM values with an arcsinh transformation. We used the fraction of variance explained (*r*^2^, also known as the coefficient of determination) to measure the predictive power of a regressor. To control for the complexity of the regressor, we used the standard adjustment 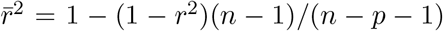, where *n* and *p* are the number of examples and regressor parameters respectively.

For the whole-gene model, we applied another linear regression model on the entire region [TSS-10kb, TTS+10kb]. This model is identical to the region-specific models, except that we computed the average feature vector over the entire region.

### 3.8 Enhancer evaluation

Following previous work [25], we evaluated an annotation according to the strength of correlation between the labels at an enhancer and the strength activity of that enhancer. We downloaded the CAGE based enhancer data from FANTOM5 website http://fantom.gsc.riken.jp/5/datafiles/latest/extra/Enhancers/human_permissive_enhancers_phase_1_and_2_expression_tpm_matrix.txt.gz. This analysis is similar to the gene expression evaluation. As the regression response value, we took the arcsinh transformation of average TPM (tags per million mapped reads) values from three replicates of GM12878 (FANTOM library ids CNhs12331, CNhs12332, and CNhs12333) as the response values. As with the gene expression evaluation, we used the average feature vector within the enhancer region as the feature vector.

### 3.9 Source code

Source code for the SSM model is available on Github at https://github.com/VertexC/State-Space-Model-on-G

## 4 Results

### 4.1 A message-passing algorithm efficiently optimizes the parameters of the SSM model

To determine whether our message-passing algorithm optimizes the parameters of the state space model as intended, we measured how the objective function of the optimization problem (regularized log likelihood, equation 4) changed over optimization iterations. We found that the message-passing algorithm quickly finds a local optimum of the objective function (Figure 2). We compared the message passing approach to an optimization algorithm that updates each position independently, and we found that the message passing approach converges much more quickly because each iteration yields a larger improvement (Figure 2).

**Figure (1).**
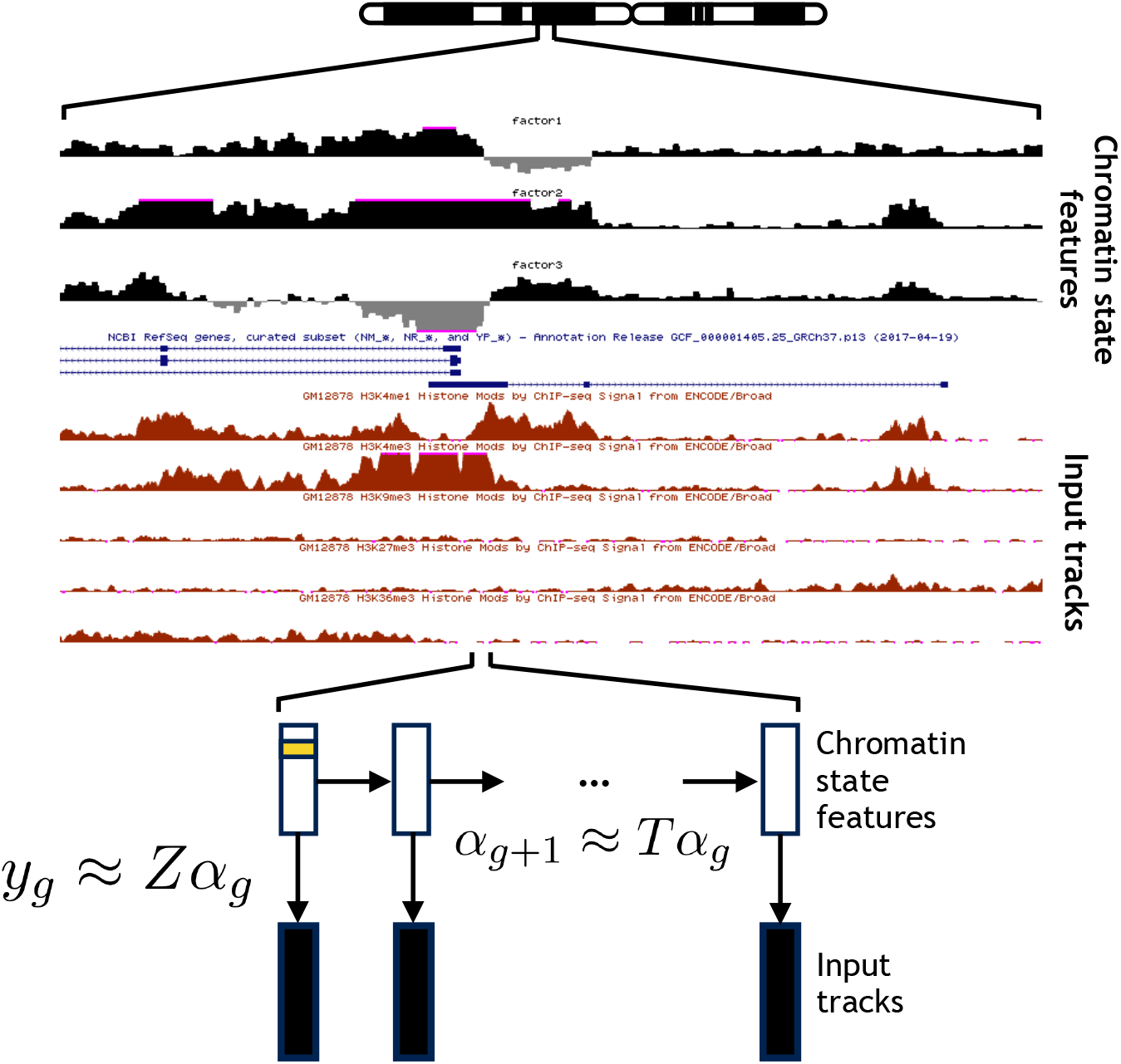
Chromatin state feature annotation. epigenome-SSM takes as input a set of genomics assays, each represented as a real-valued track over the genome. It outputs a set of realvalued chromatin state features for each position in the genome, using a Kalman filter state space model.

**Figure (2).**
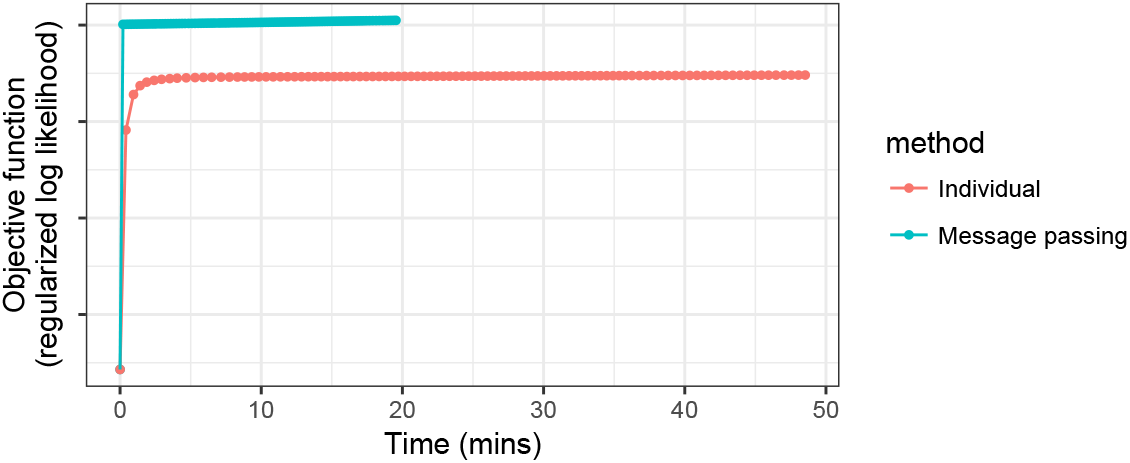
SSM model optimization. Vertical axis indicates the SSM model’s objective function. Horizontal axis indicates running time. Performed on a simulated genome of length 30k bp, with 100-iteration optimization.

### 4.2 Chromatin state features at promoters are predictive of gene expression

To evaluate annotation methods, we compared annotations to RNA-seq gene expression data, following previous work [24, 16] (Section 3.7). Briefly, in a high-quality annotation, highly-expressed genes should be annotated in a distinct way from the rest of the genome. In other words, there should be a strong correlation between the annotation at the promoter and the gene’s expression level, as measured by RNA-seq. Specifically, we evaluated the quality of an annotation according to the variance explained by a predictive model that takes as input the annotation state at a gene’s promoter and outputs predicted RNA-seq expression.

We found that all models are predictive of gene expression, but the SSM model outperforms alternatives by this measure (Figure 3a). An SSM annotation explains significantly more variance (Adj *r*^2^ = 0.17 for *k* = 3) than a discrete HMM, whether or not the HMM models continuous input with a Gaussian distribution or models discrete input with a Bernoulli distribution (0.09 and 0.13 respectively). Part of this improvement is a result of the greater richness of a continuous model—if we produce a continuous annotation from the HMM model by using the probability of each state, the performance improves (0.11 and 0.13 for the continuous and discrete input models respectively). However, this performance is still worse than the SSM annotation, indicating the importance of using a model that is intrinsically continuous. Continuous PCA and NMF models perform worse than SSM in the whole-gene model (0.11 and 0.13 respectively), indicating the value of using a model that incorporates the genomic axis (although PCA and NMF perform slightly better in the region-based regression).

**Figure (3).**
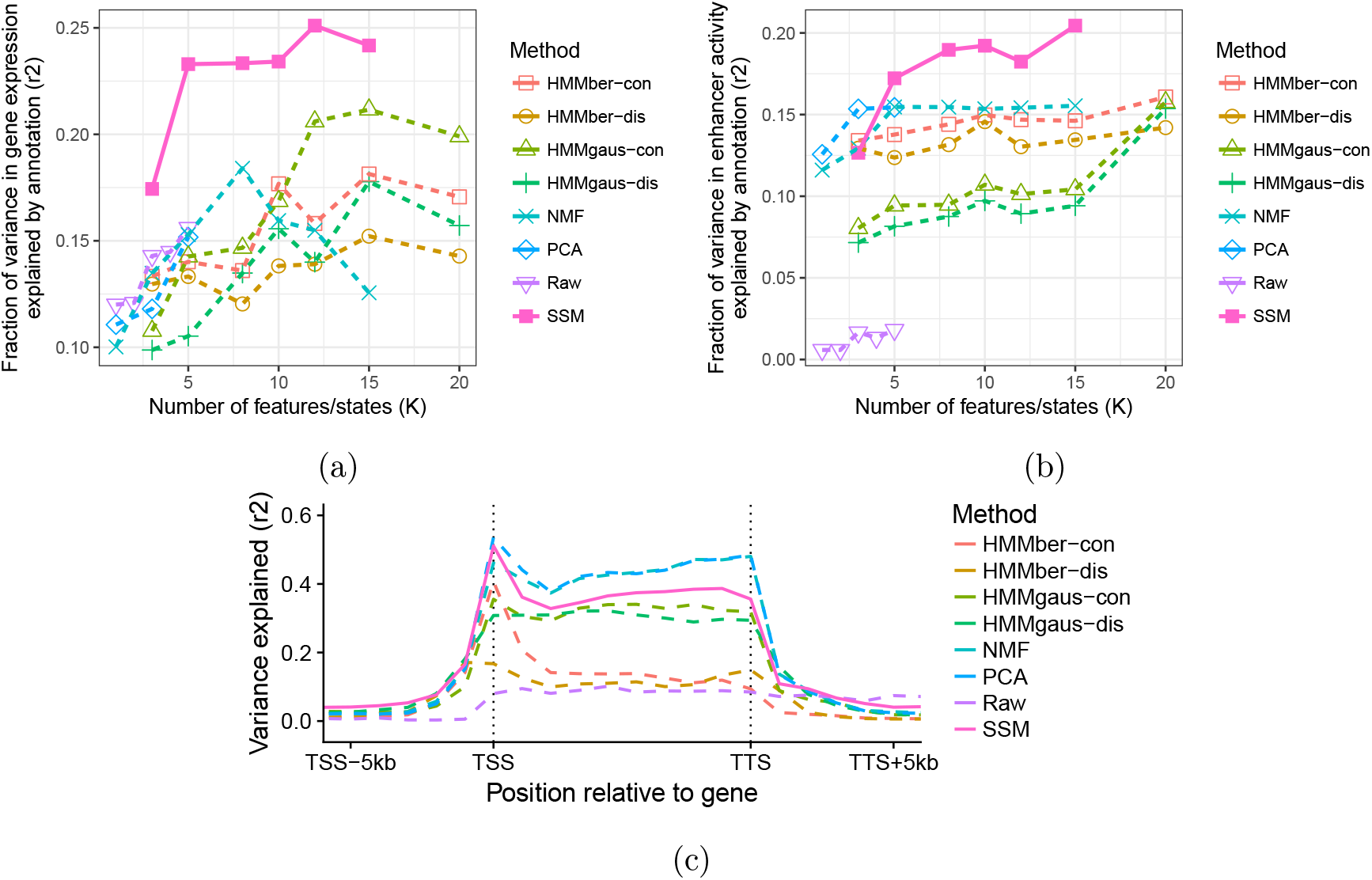
Evaluation of annotations relative to gene expression and enhancer activity. (a) Vertical axis is the fraction of variance in gene expression explained (*r*^2^, Section 3.7). Horizontal axis is the number of features or states in a given continuous or discrete model respectively (*k*). Each line corresponds to a model. (b) Same as (a), but using enhancer activity from FANTOM5 (Section 3.8). (c) Same as (a), but using the annotation at a specific relative to the gene rather than the entire TSS as the predictor (Section 3.7). The horizontal axis indicates which region relative to the gene was used as a predictor. All methods use *k* = 3.

Moreover, the superior performance of the SSM is maintained even when the HMM model uses more labels than the number of SSM features. Even for the largest number of labels we tried (*k* = 20), the performance of any HMM model was no higher than *r*^2^ = 0.21, which is lower than the performance of the SSM model with 5 features (0.23). This comparison offsets the potential disadvantage that continuous features are more complex than discrete labels. If one is interested in obtaining a very simple annotation, according to this analysis, it is preferable to use a continuous model with a small number of features rather than a discrete model with many labels.

We found that different types of annotations differed in which positions relative to the transcription start site were predictive of expression (Figure 3c). As we expected, positions near the transcription start site (TSS) are most predictive of expression, whereas annotations at positions over 3 kp from the TSS have less predictive power. Notably, however, discrete annotations at the TSS itself are no more predictive of expression than annotations at its flanks. This is likely because the discrete methods label the TSS as such regardless of whether it is expressed, whereas the continuous features can express both whether or not the position is a TSS and whether or not it is expressed. This behaviour shows how a continuous model can incorporate longer-range dependencies between positions.

### 4.3 Chromatin state features at enhancer elements are predictive of enhancer activity

We further evaluated these annotation methods by measuring how predictive each annotation is of experimentally-validated enhancer elements, again following previous work [24]. We used FANTOM5 enhancer RNA data as a measurement of the activity of each enhancer element. Although the SSM model performs slightly worse with *k* = 3, it performs significantly better (Adj *r*^2^ = 0.17, *k* = 5) than all alternatives for *k* ≥ 5 (0.01-0.15).

### 4.4 Chromatin state features recapitulate known genome biology

While the results above show quantitatively that chromatin state features are predictive of many genomic phenomena, we additionally found that these features qualitatively recapitulate known genome biology. We focus here on an annotation using three features (*k*=3).

Feature 2 is a general mark of activity, indicating presence of all histone modifications except for the heterochromatic mark H3K9me3 (Figure 4a). Thus, both promoters and enhancers are characterized by feature 2 (Figures 4d and 4f). In fact, transcription termination sites (TTSs) are also enriched for feature 2, indicating previously-reported [9] but under-appreciated regulatory state around TTSs (Figure 4e). Feature 2 is strongly predictive of gene expression; most promoters with large positive values of feature 2 are expressed, while most without feature 2 (with low value) are not expressed (Figure 4c).

**Figure (4).**
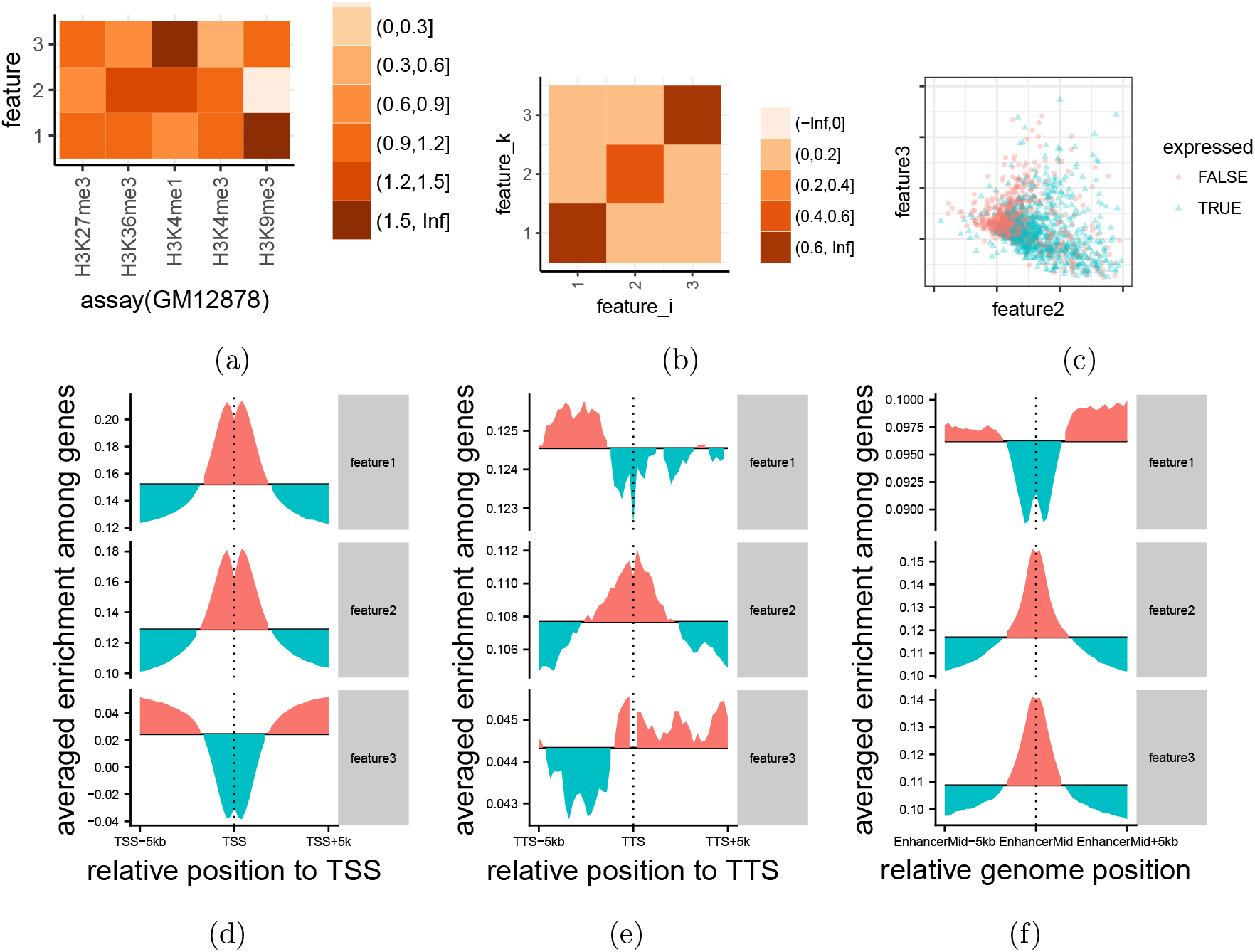
(a) **Relationship of features to the input genome data**. Color corresponds to the mean signal value of a given assay at positions annotated with a given feature. (b) **Relationship between features in neighboring positions**. Color in cell *i,k* represents the correlation of feature *i* at one position with feature *k* in the following position. (c) **Distribution of promoter features**. Each point corresponds to a gene. Teal points correspond to genes in the top 33% of gene expression; orange points correspond to genes in the lower 66%. Horizontal and vertical axes indicate average values of feature 2 and feature 3 respectively at the gene’s promoter (average value in the 10 kb upstream of the TSS). For the sake of visibility, the plot shows a random 10% subset of all genes. (d-f) **Enrichment of chromatin state features relative to genomic elements**. Vertical axis indicates average feature value. (d,e) Transcription start sites (TSSs) and transcription termination sites (TTSs). (f) Enhancers.

Feature 3 is mark of enhancer-specific activity. It indicates the presence of the enhancer-associated mark H3K4me1, but not the promoter-associated mark H3K4me3 (Figure 4a). Thus, enhancers are associated with large positive values of feature 3 (Figure 4d), while promoters are associated with large negative values of this feature (Appendix Figures 5,6).

**Figure (5).**
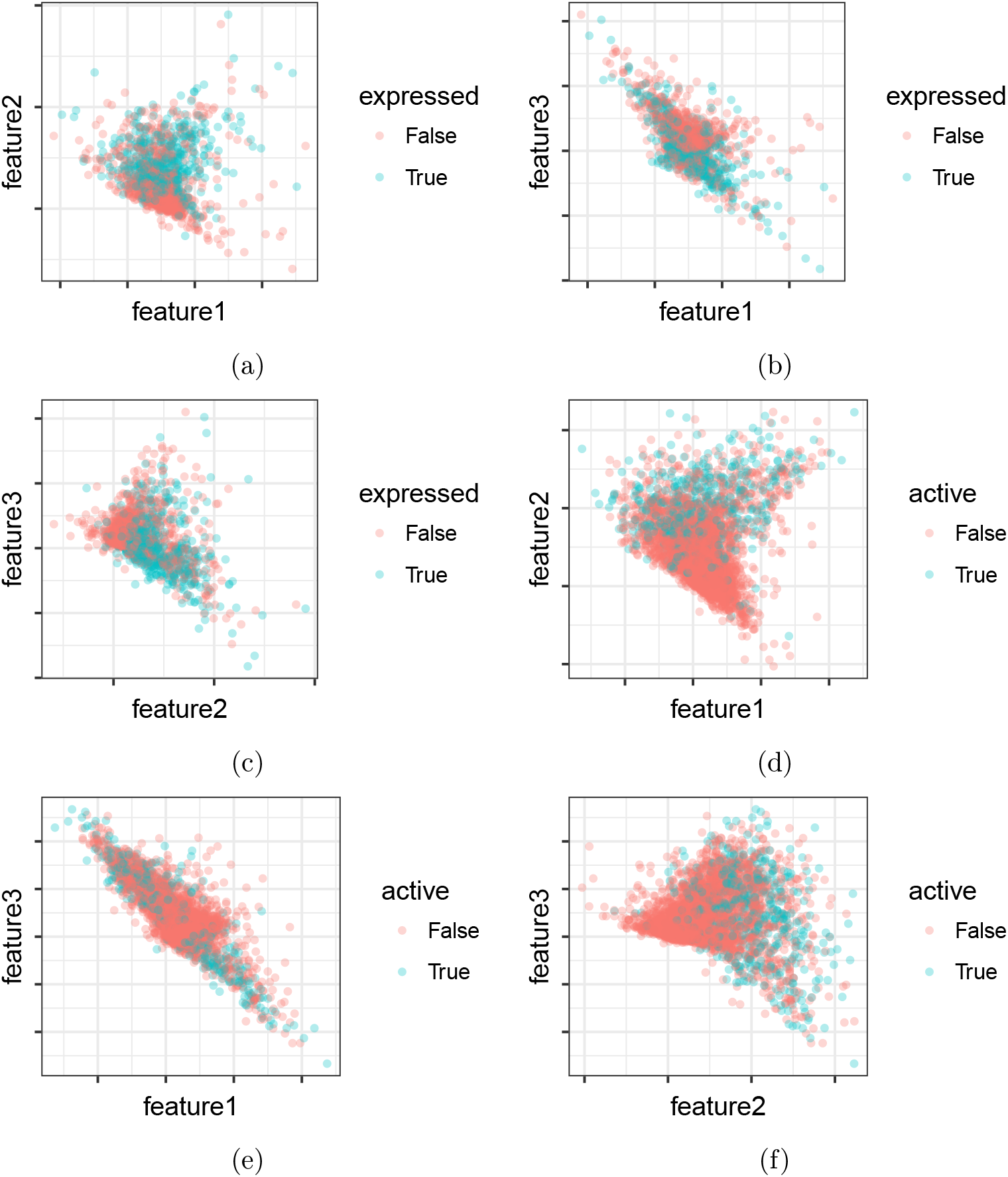
(a-c) **Distribution of promoter features.** Each point corresponds to a gene. Teal points correspond to genes in the top 33% of gene expression; orange points correspond to genes in the lower 66%. Horizontal and vertical axes indicate average values of two features respectively at the gene’s promoter (average value in the 10 kb upstream of the TSS). For the sake of visibility, the plot shows a random 10% subset of all genes. (d-f) **Distribution of enhancer features.** Same as (a-c), but expression corresponds to CAGE enhancer activity.

**Figure (6).**
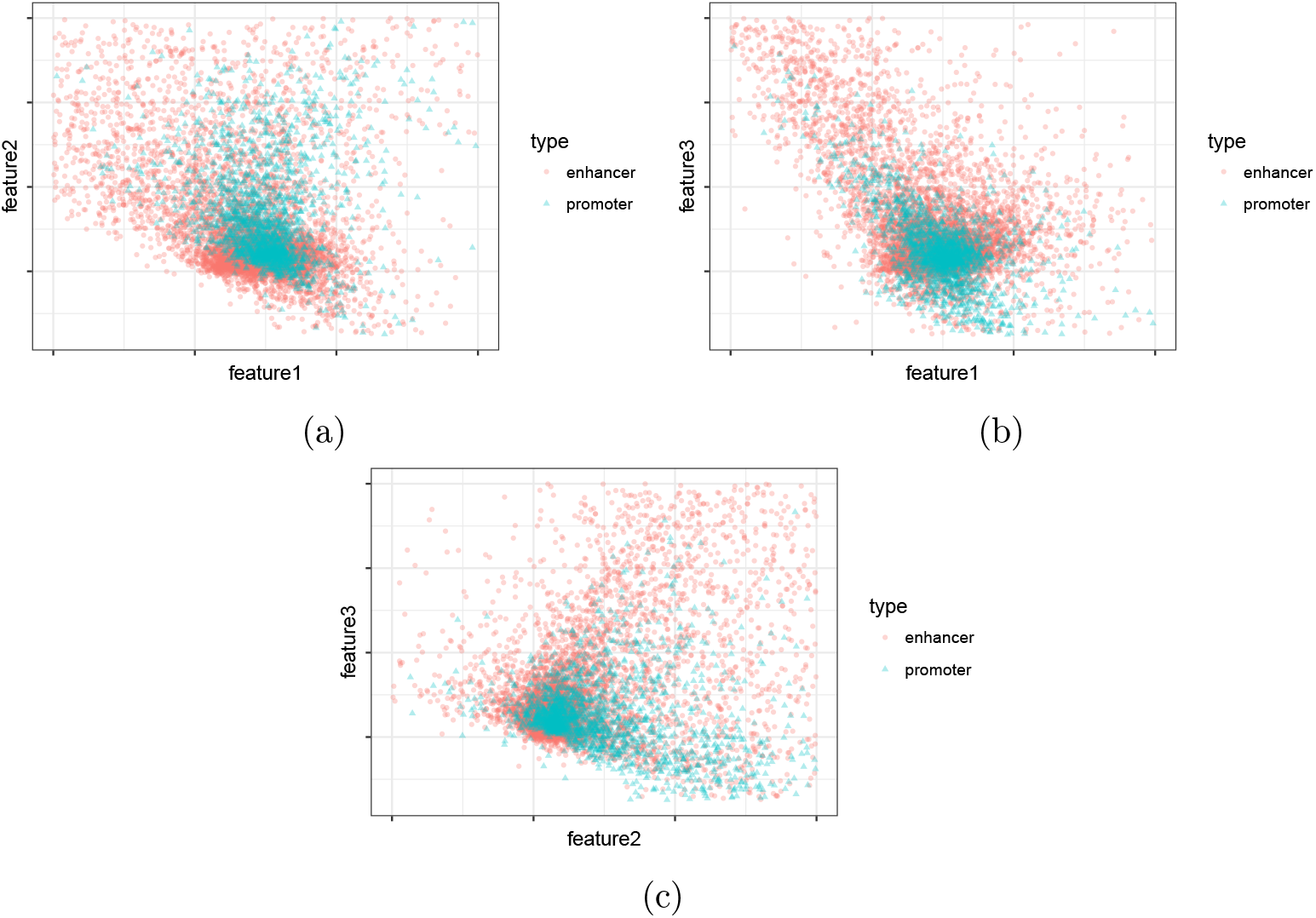
Distribution of promoter and enhancer features.

Feature 1 is a general mark of broad activity, both broad transcribed regions marked by H3K36me3 as well as broad repressed regions marked by H3K9me3 and H3K27me3 (Figure 4a). Whether a given broad region is active or repressed is indicated by feature 2, which has opposite values for these two categories of marks.

## 5 Discussion

In this work, we introduced continuous chromatin state features for genome annotation. These chromatin state features are analogous to existing discrete chromatin state labels, but continuous features have several benefits: they can represent varying strength among elements, and they can easily represent combinatorial patterns of activity. Due to these benefits, we showed that chromatin state features outperform existing discrete annotations at predicting gene expression and enhancer activity. While continuous features are somewhat more complicated to interpret than discrete labels, we showed that a small number of continuous features outperform even a large number of discrete labels in all of our evaluations. Moreover, chromatin state easily lend themselves to expressive visualizations. Because continuous annotations maintain much more of the information in the input data than discrete annotations do, they are more useful for complex downstream applications. For example, a variant effect predictor might take chromatin state features as input in order to predict the functional impact of a given mutation. This is preferable to using raw tracks for two reasons. First, a small number of chromatin state features concisely summarize a large number of input tracks and therefore a predictive model based on these features will be less prone to overfitting. Second, chromatin state features can be used for variant effect interpretation; that is, a model could report that its prediction of high variant effect is due to the fact that a specific feature is present at that position. Such interpretation is more difficult with raw tracks because most types of activity are associated with a combination of many marks.

In the future, we plan to apply this approach to create chromatin state features for all tissues with sufficient available data. Also, there are many modifications that may improve the base SSM method. In addition to the modifications that have proved successful for HMM-based models such as handling missing data, the interpretability of chromatin state feature annotations may be improved by encouraging sparsity, smoothness or restricting values to be positive.

## 6 Appendix

### 6.1 pseudo code of SSM

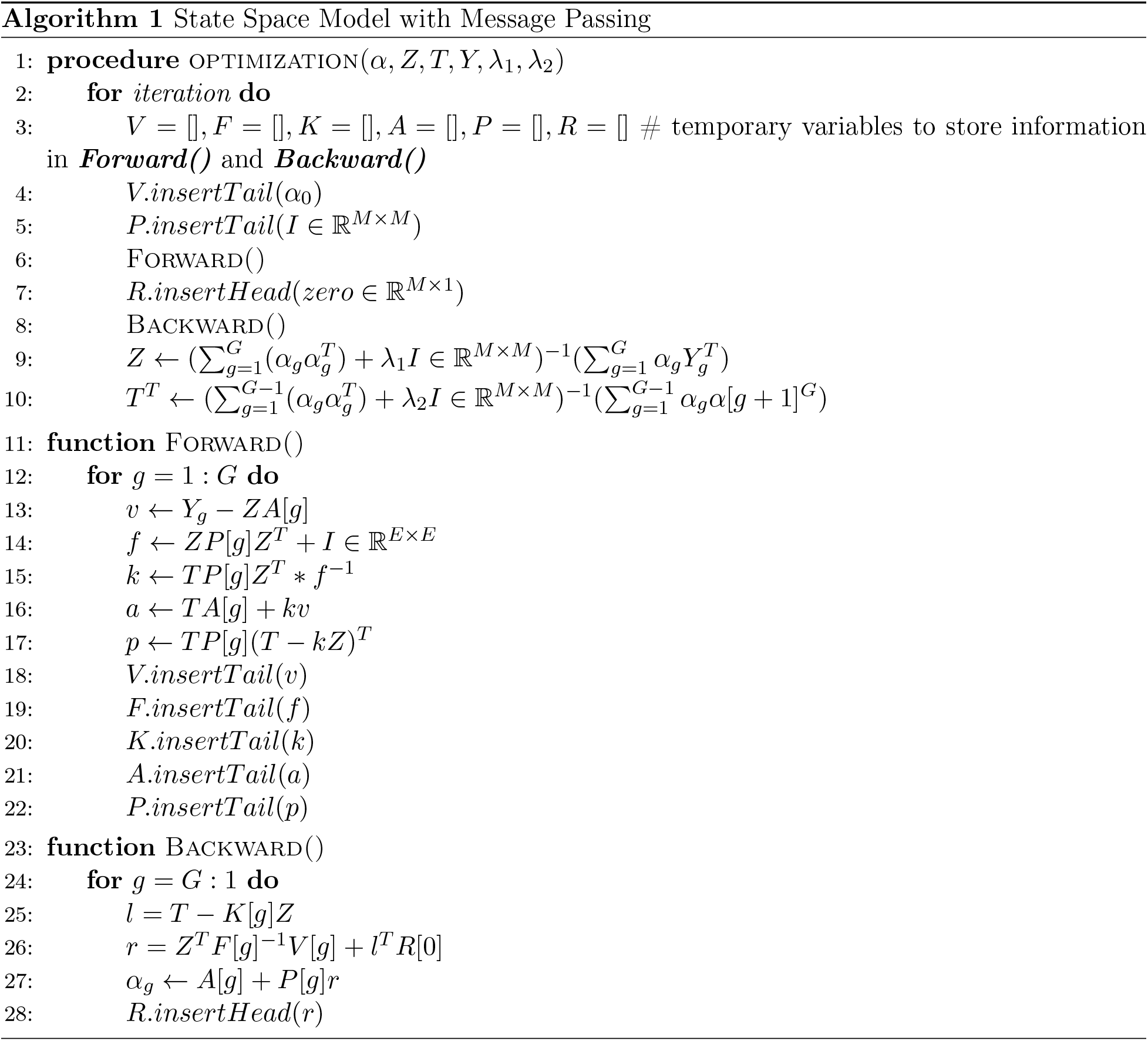

